# NEK10 tyrosine phosphorylates β-catenin to regulate its cytoplasmic turnover

**DOI:** 10.1101/2022.12.23.521717

**Authors:** Previn Dutt, Nasir Haider, Samar Mouaaz, Lauren Podmore, Vuk Stambolic

## Abstract

Nek kinases are involved in regulating several different elements of the centrosomal cycle, primary cilia function, and DNA damage responses. Unlike the other members of the Nek family, which are serine-threonine kinases, Nek10 preferentially targets tyrosines. Nek10 appears to have a broad role in DNA damage responses, regulating a MAPK-activated G2/M checkpoint following UV irradiation and influencing the p53-mediated activation induced by genotoxicity. In an attempt to identify additional Nek10 functions, we characterized the effect of Nek10 deletion in lung cancer cells, where it is relatively highly expressed. Nek10 absence led to an increase in both the signaling and adherens junctions pools of β-catenin. Mechanistically, Nek10 associates with the Axin complex where it phosphorylates β-catenin at Tyr30, located within the regulatory region governing β-catenin turnover. In the absence of Nek10 phosphorylation, GSK3-mediated phosphorylation of β-catenin, a prerequisite for its turnover, was significantly impaired. Stabilization of β-catenin driven by Nek10 loss diminished the ability of cells to form tumorspheres in suspension, grow in soft agar, and colonize mouse lung tissue following tail vein injections.

## Introduction

Nek10 is one of eleven members of the family of human Nek kinases implicated in regulation of various aspects of centrosome biology, primary cilia function, and DNA damage responses. While Nek2 and Nek5 are primarily involved in centrosome disjunction, a signaling module consisting of Nek6, Nek7, and Nek9 impacts several different mitotic functions, including centrosome positioning, spindle assembly and cytokinesis (Fry et al. 2017). Beyond these mitotic roles, Nek1 and Nek8, in particular, have been linked to ciliopathies such as polycystic kidney disease (Moniz et al., 2011a, de Oliveira et al., 2020).

Whereas the other members of the Nek family are serine-threonine kinases, Oriented Peptide Library Screening revealed that Nek10 has a preference for phosphorylation of tyrosine residues harboring phenylalanine, tryptophan, histidine, or leucine at the +1 position (van de Kooij et al., 2019). Nek10 has been reported to interact directly with the MAP kinase signaling module and promote phosphorylation of ERK1/2 (Moniz et al., 2011b). While Nek10 is dispensable for ERK activation in response to mitogenic factors, it augments MAP kinase throughput related to a G2/M checkpoint engagement following UV irradiation. Most recently, it has been demonstrated that Nek10 also affects the p53-mediated response to DNA damage by phosphorylating p53 on Tyr327 (Haider et al., 2020).

Work presented here implicates Nek10 in the regulation of β-catenin levels which are exquisitely controlled in cells. While the majority (∼90%) of β-catenin is relatively stable and associated with cadherins in adherens junctions at the cell membrane, the remaining β-catenin is found in a highly dynamic signaling pool that is rapidly turned over in the absence of Wnt pathway signaling input (Gerlach et al., 2014). This turnover involves the recruitment of free cytoplasmic β-catenin into the Axin complex, followed by its sequential phosphorylation by CK1 at Ser45, and then by GSK3 at Ser33, Ser37, and Thr41 (Valenta et al., 2012; Verheyen and Gottardi, 2010). These phosphorylation events promote recruitment of the β-Trcp ubiquitin E3 ligase, resulting in the ubiquitination and proteosomal degradation of β-catenin (van Kappel and Maurice, 2017). APC is an indispensable component of the turnover complex, though its precise function and, more broadly, the mechanistic steps by which phosphorylated β-catenin is ubiquitinated and degraded remain unclear. Upstream signaling triggered by the binding of Wnt ligands to Frizzled receptors disrupts degradation of β-catenin, leading to its accumulation in the nucleus where it interacts with the TCF/LEF transcriptional co-factors to activate target genes (Nusse and Clevers, 2017).

Wnt-β-catenin signaling has well established roles in controlling body patterning during embryogenesis and development, as well as regulating self-renewal and differentiation during adult tissue homeostasis (Pond et al., 2020; Steinhart and Angers, 2018). Wnt morphogens guide cell fate determination in target cells by promoting the appropriate spatio-temporal transcriptional programs. Fine tuning of Wnt-mediated signaling is achieved through the concerted action of additional secreted factors, such as BMPs (bone morphogenetic proteins), as well as intracellular modifiers, acting either through the β-catenin pathway itself, or distinct developmental pathways capable of communicating with β-catenin throughput. More recently, the role of Wnt-β-catenin in regulating the stem cell niche in adult tissues, particularly the intestine, bone, and skin, has become better understood. For instance, a Wnt signaling gradient has been observed within intestinal crypts whereby Wnt ligands produced from underlying myoepithelial cells results in higher expression of Wnt target genes at the base of the crypt.

Deregulation of the Wnt/β-catenin pathway is common in a variety of cancers (Jackstadt et al., 2020). The archetypal β-catenin-driven oncogenic program is found in colorectal cancers, approximately 70% of which feature loss of function of the APC gene, resulting in elevated β-catenin levels. Roughly 50% of hepatocellular carcinomas harbor various activating mutations targeting the Wnt pathway, with half of these found in the β-catenin gene (Khalaf et al., 2018). While upregulation of β-catenin does not by itself induce lung cancer, it has been shown to dramatically affect the progression of tumors driven by other oncogenic programs, such as KRAS and EGFR activation (Pacheco-Pinedo et al., 2011; Nakayama et al., 2014). A meta-analysis of non-small cell lung cancer (NSCLC) studies concluded that reduced membrane β-catenin and elevated nuclear β-catenin were associated with poor clinical outcome, although no significant overall correlation between abnormal β-catenin levels and prognosis could be confirmed (Jin et al., 2017). Additionally, Wnt target gene expression has been reported to contribute to lung adenocarcinoma metastasis (Nguyen et al., 2009).

Here we report that Nek10 deficiency in A549 lung cancer cells results in an increase in β-catenin levels, both in the signaling and adherens junction pools. Elevated β-catenin levels are a consequence of a reduction in Nek10-mediated phosphorylation at Tyr30 within the N-terminal regulatory region of the protein that governs its turnover. Reduced β-catenin Tyr30 phosphorylation, in the absence of Nek10, impaired GSK3-mediated phosphorylation events that are a prerequisite for β-catenin turnover and subsequent recruitment of the β-Trcp E3 ubiquitin ligase. At a cellular level, stabilization of β-catenin diminished the ability of A549 cells to form tumorspheres in suspension, grow in soft agar, and colonize mouse lung tissue following tail vein injections.

## Results

### Nek10 deletion in the A549 lung cancer line leads to elevated cellular levels of β-catenin

The Nek10 kinase is relatively highly expressed in both mouse and human lung tissue (Fig.S1). Lower Nek10 expression correlates with better clinical outcomes in patients diagnosed with lung adenocarcinomas (Fig.S2). To evaluate potential Nek10 functions in lung cancer biology, we deleted a Nek10 kinase domain exon encoding the critical DFG motif in A549 cells, as described previously (Haider et al., 2020). Initial characterization of various signaling pathways within the resulting knockout cell lines indicated that β-catenin was significantly upregulated in multiple clonal lines (Fig.1A, Fig.S3A). Cell fractionation revealed that β-catenin was elevated in both the nuclear and cytosolic fractions of *NEK10*^*Δ/Δ*^ A549 cells (Fig.1B). GST-ICAT, a reagent designed to specifically capture the signaling pool of β-catenin (Flozak et al., 2016), pulled down more β-catenin from Nek10-null than wild-type cell lysates (Fig.1C). In *NEK10*^*Δ/Δ*^ cells, more β-catenin co-immunoprecipitated with E-cadherin than in their wild-type counterparts (Fig.1D), and an increase in β-catenin immunostaining was observed at the cell membranes, where roughly 90% of β-catenin is typically localized within adherens junctions (Fig.S3B). Elevated β-catenin in *NEK10*^*Δ/Δ*^ cells was not a consequence of hyperactivation of the Wnt pathway, as phosphorylation of the Lrp6 co-receptor under both basal and Wnt-3a-induced conditions was indistinguishable in Nek10-proficient and -deficient cell lines (Fig.S4A). Quantitative RT-PCR analysis revealed no significant changes in β-catenin expression levels or upregulation in the expression levels of the known β-catenin target gene, Axin2 (Fig.S4B).

**Figure 1.**
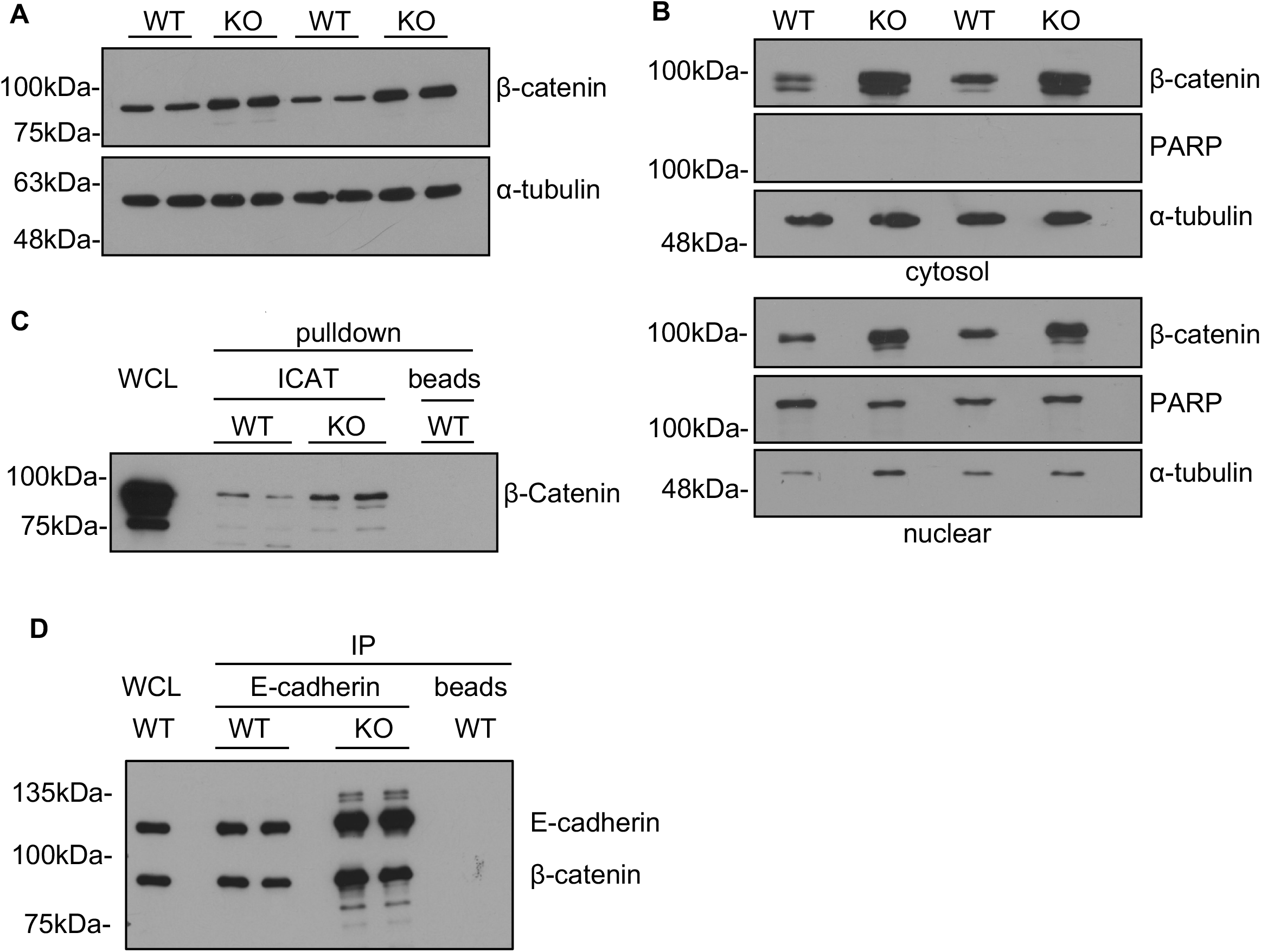
Nek10 deletion in the A549 lung cancer line results in elevated β-catenin in both the signaling and adherens junction pools. (A) Total β-catenin levels in whole cell lysates of *NEK10*^*+/+*^ and *NEK10*^*Δ/Δ*^ A549 lines. α-tubulin was used as a loading control. (B) β-catenin levels in the cytosolic and nuclear fractions of *NEK10*^*+/+*^ and *NEK10*^*Δ/Δ*^ cells. α-tubulin and PARP levels were used as markers of the cytosolic and nuclear fractions, respectively. (C) Total β-catenin pulled down by GST-ICAT from *NEK10*^*+/+*^ and *NEK10*^*Δ/Δ*^ A549 lysates. (D) The levels of β-catenin co-immunoprecipitating with E-cadherin in *NEK10*^*+/+*^ and *NEK10*^*Δ/Δ*^ A549 cells.

### Nek10 influences GSK3 phosphorylation of β-catenin by specifically targeting β-catenin at tyrosine 30

β-catenin turnover within the Axin destruction complex involves its phosphorylation by CK1 at Ser45, which primes for further GSK3β phosphorylation at Ser33, Ser37, and Thr41 (Fig.2A) (Nusse and Clevers, 2017; van Kappel and Maurice, 2017). These sequential modifications, in turn, promote ubiquitination and proteosomal degradation of β-catenin. The turnover of free β-catenin is highly dynamic, such that the aforementioned phosphorylated species are present at limited levels in cycling cells. When wild-type and *NEK10*^*Δ/Δ*^ A549 cells were treated with MG132, to arrest proteosomal turnover, *NEK10*^*Δ/Δ*^ cells displayed increased CK1-phosphorylated β-catenin compared to their wild-type counterparts, consistent with the elevated total β-catenin levels (Fig.2B). In contrast, GSK3β-phosphorylated β-catenin levels were reduced in *NEK10*^*Δ/Δ*^ cells (Fig.2B). This was not the result of an overall change in the GSK3 activity, as determined by comparing the levels of GSK3 phosphorylation of Tau in *NEK10*^*+/+*^ *and NEK10*^*Δ/Δ*^ cells (Fig.S5).

**Figure 2.**
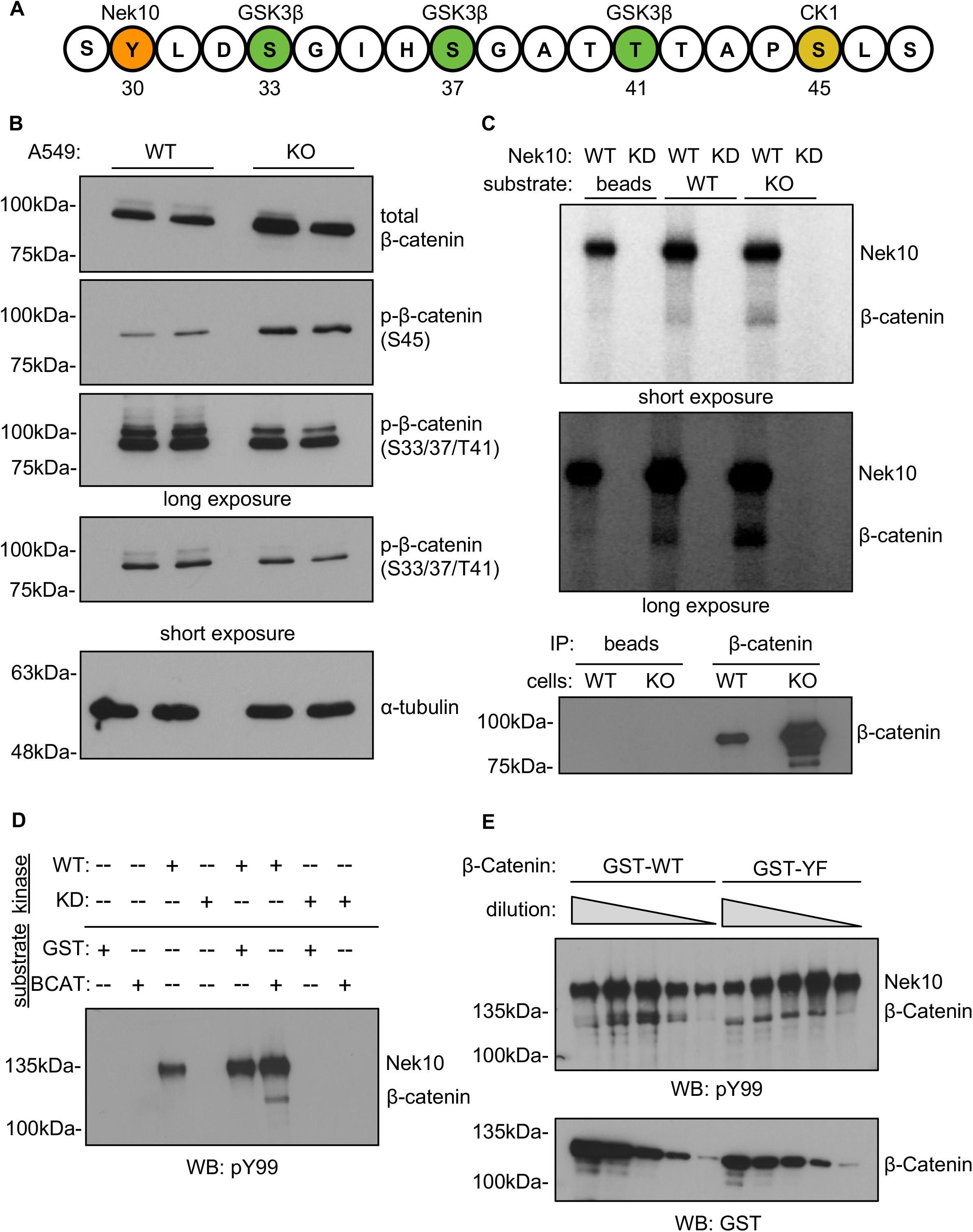
Nek10 influences phosphorylation of the critical β-catenin regulatory sites and specifically targets β-catenin at tyrosine 30. (A) Schematic of the locations of the critical N-terminal CK1 and GSK3 phosphorylation sites controlling β-catenin turnover, in relation to a putative Nek10-targeted site. (B) β-catenin phosphorylation at Ser45 and Ser33/Ser37/Thr41, relative to total β-catenin levels, in *NEK10*^*+/+*^ and *NEK10*^*Δ/Δ*^ A549 cells. α-tubulin was used as a loading control. (C) A radioactive kinase assay performed using recombinant wild-type (WT) or kinase-dead (KD) Nek10 to phosphorylate β-catenin from *NEK10*^*+/+*^ and *NEK10*^*Δ/Δ*^ A549 cells. (D) A non-radioactive kinase assay testing the ability of recombinant wild-type (WT) or kinase dead (KD) Nek10 to tyrosine phosphorylate recombinant GST or GST-β-catenin (BCAT). (E) A non-radioactive kinase assay comparing the ability of recombinant wild-type Nek10 to tyrosine phosphorylate recombinant wild-type (WT) and mutant (Y30F) GST-β-catenin. In the mutant β-catenin, the putative Nek10 tyrosine target was changed to a non-phosphorylatable phenylalanine. A substrate dilution series was used to facilitate comparison.

Although by sequence homology a member of the Nek family of serine-threonine kinases, Nek10 preferentially phosphorylates tyrosines, with phenylalanine, tryptophan, leucine or histidine at the Tyr+1 position (van de Kooij et al., 2019). The β-catenin sequence contains a Tyr-Leu motif at Tyr30 which is proximal to the aforementioned CK1- and GSK3β-phosphorylation sites, as well as the subsequent ubiquitination sites at Lys19 and Lys49 (Fig.2A). In an in vitro ^32^P kinase assay, Nek10 phosphorylated β-catenin immunopurified from both *NEK10*^*+/+*^ and *NEK10*^*Δ/Δ*^ A549 cells, with the level of phosphorylation corresponding to the relative amounts of β-catenin in the immunoprecipitates (Fig.2C). Nek10 phosphorylated tyrosines in β-catenin, judged by a non-radioactive kinase assay using recombinant GST-β-catenin as substrate and pY99 anti-tyrosine antibody immunoblotting as readout (Fig.2D). In the same assay, Nek10 phosphorylated the GST-β-catenin Y30F mutant to a lesser degree, suggesting that Tyr30 is a Nek10 phosphorylation target in β-catenin (Fig.2E).

### Loss of β-catenin tyrosine 30 phosphorylation by Nek10 impedes its GSK3-mediated phosphorylation, leading to a rise of intracellular β-catenin levels

To assess the effect of the β-catenin Tyr30 phosphorylation in cells, we expressed N-terminal HA-tagged wild-type and Y30F mutant β-catenin in HEK293T cells. Y30F β-catenin protein presented at considerably higher levels compared to the wild-type, suggestive of increased stability of the mutant (Fig.3A). The exogeneous β-catenin appropriately localized to both the (ICAT-associated) signaling and (E-cadherin-associated) adherens junction pools, with the mutant protein elevated in both (Fig.S6A-B). A similar effect was noted with C-terminal Myc-tagged β-catenin bearing the Y30F mutation (Fig.S6C). Moreover, when the tagged β-catenin proteins were immunoprecipitated, the Y30F mutant exhibited less relative phosphorylation at GSK3β-target sites compared to the wild-type protein (Fig.3B) similar to the effect observed in Nek10-null A549 cells (see Fig.2B).

**Figure 3.**
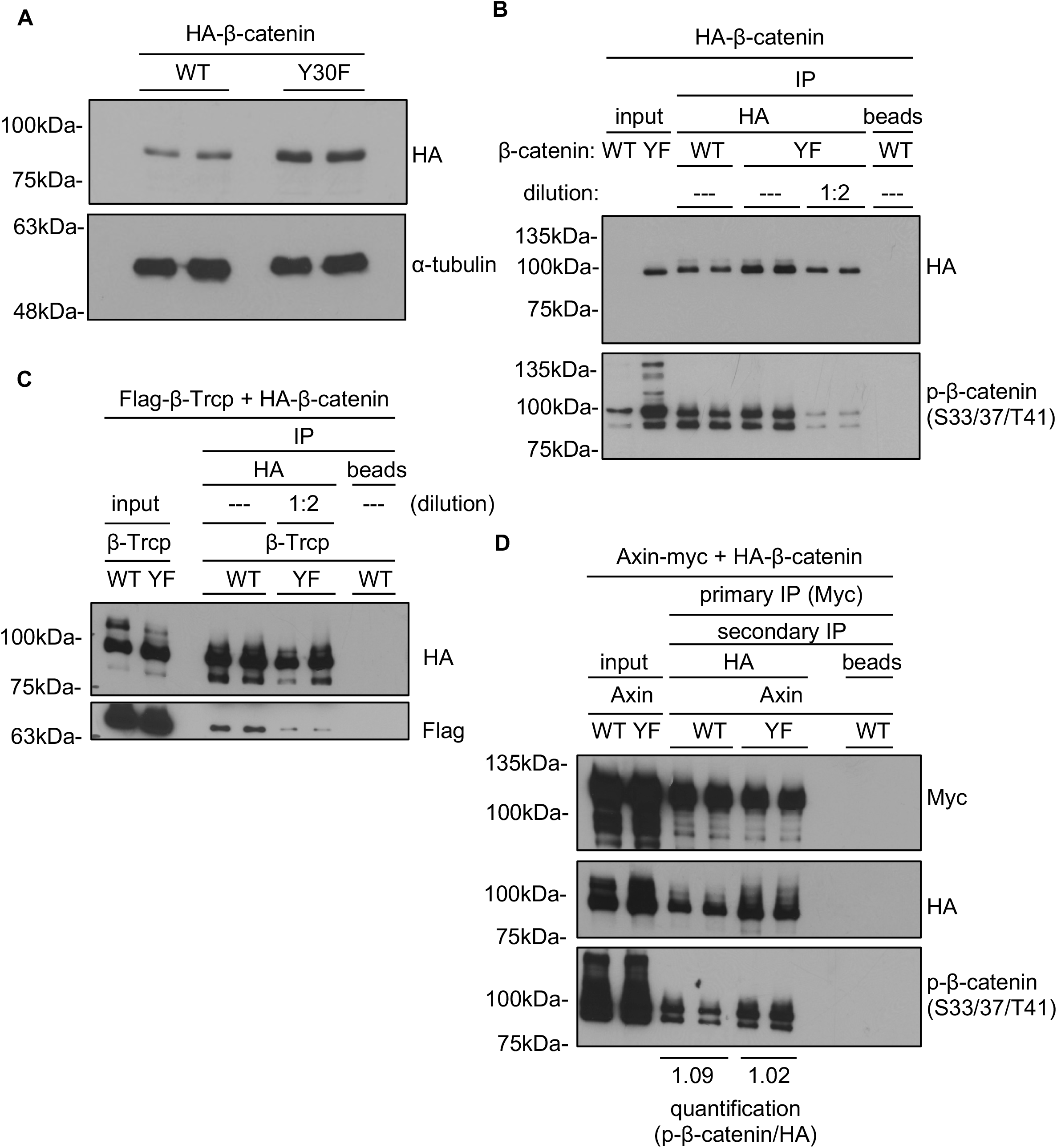
Loss of Nek10 phosphorylation at tyrosine 30 impedes GSK3β mediated-phosphorylation of β-catenin and increases its cellular levels. (A) The levels of HA-tagged wild-type (WT) and mutant (Y30F) β-catenin exogeneously expressed in HEK293T cells. (B) The levels of Ser33/Ser37/Thr41 phosphorylation in immunoprecipitated HA-tagged wild-type (WT) and mutant (Y30F) β-catenin. Since the mutant protein is expressed at higher levels, phospho-antibody signals were normalized to the levels of total β-catenin in the immunoprecipitates. (C) Comparison of Flag-tagged β-Trcp co-immunoprecipitating with HA-tagged wild-type (WT) and mutant (Y30F) β-catenin. The immunoprecipitated mutant β-catenin was diluted to facilitate comparison with the wild-type protein. (D) The amount of total and Ser33/Ser37/Thr41 phosphorylated HA-tagged wild-type and mutant (Y30F) β-catenin co-immunoprecipitating with Myc-tagged Axin. To separate HA-tagged β-catenin from the endogeneous protein, Myc-Axin immunoprecipitates (primary IP) were first eluted and then re-immunoprecipitated with anti-HA (secondary IP).

To determine if the differential phosphorylation impacted β-catenin turnover, we assessed association of β-catenin with its cognate E3 Ubiquitin ligase, β-Trcp, and found that the Y30F mutant displayed reduced β-Trcp engagement (Fig.3C). Interestingly, a greater association of total and GSK3β-phosphorylated β-catenin with the Axin complex was observed in HEK293T cells expressing exogeneous Y30F β-catenin, roughly correlating with the greater levels of β-catenin available for entry into the Axin complex (Fig.3D).

### β-catenin Y30F mutant is responsive to upstream Wnt signaling, GSK3 inhibition, and undergoes proteosomal degradation

Cellular β-catenin levels increase following treatment with either Wnt-3a, LiCl, or MG132, all of which impede the degradation of the protein, albeit at different points in the process (Li et al., 2012; van Kappel and Maurice, 2017). Treatment of HEK293T cells with Wnt-3a resulted in stabilization of HA-tagged β-catenin (Fig.4A). This was true for both wild-type and Y30F β-catenin, demonstrating that the mutant protein retained responsiveness to upstream signaling. While the mechanisms by which Wnt stabilizes β-catenin are still not completely agreed upon, the Y30F mutation contributes additional stabilization beyond that achieved by Wnt alone. A similar effect was observed when cells treated with LiCl, a GSK3 inhibitor, or the proteosomal inhibitor, MG132, both of which led to elevation of cellular levels of both wild-type and Y30F β-catenin, with the mutant protein accumulating at higher levels (Fig.4B-C). In order to separate the responsive signaling pool from the more stable adherens junction pool, parallel analyses were carried out looking specifically at the levels of HA-tagged β-catenin co-immunoprecipitating with Flag-ICAT in the presence of Wnt-3a, LiCl, and MG132 (Fig.4D-F). Each treatment induced a buildup in both wild-type and mutant β-catenin within the signaling pool, with the mutant protein stabilizing at higher levels.

**Figure 4.**
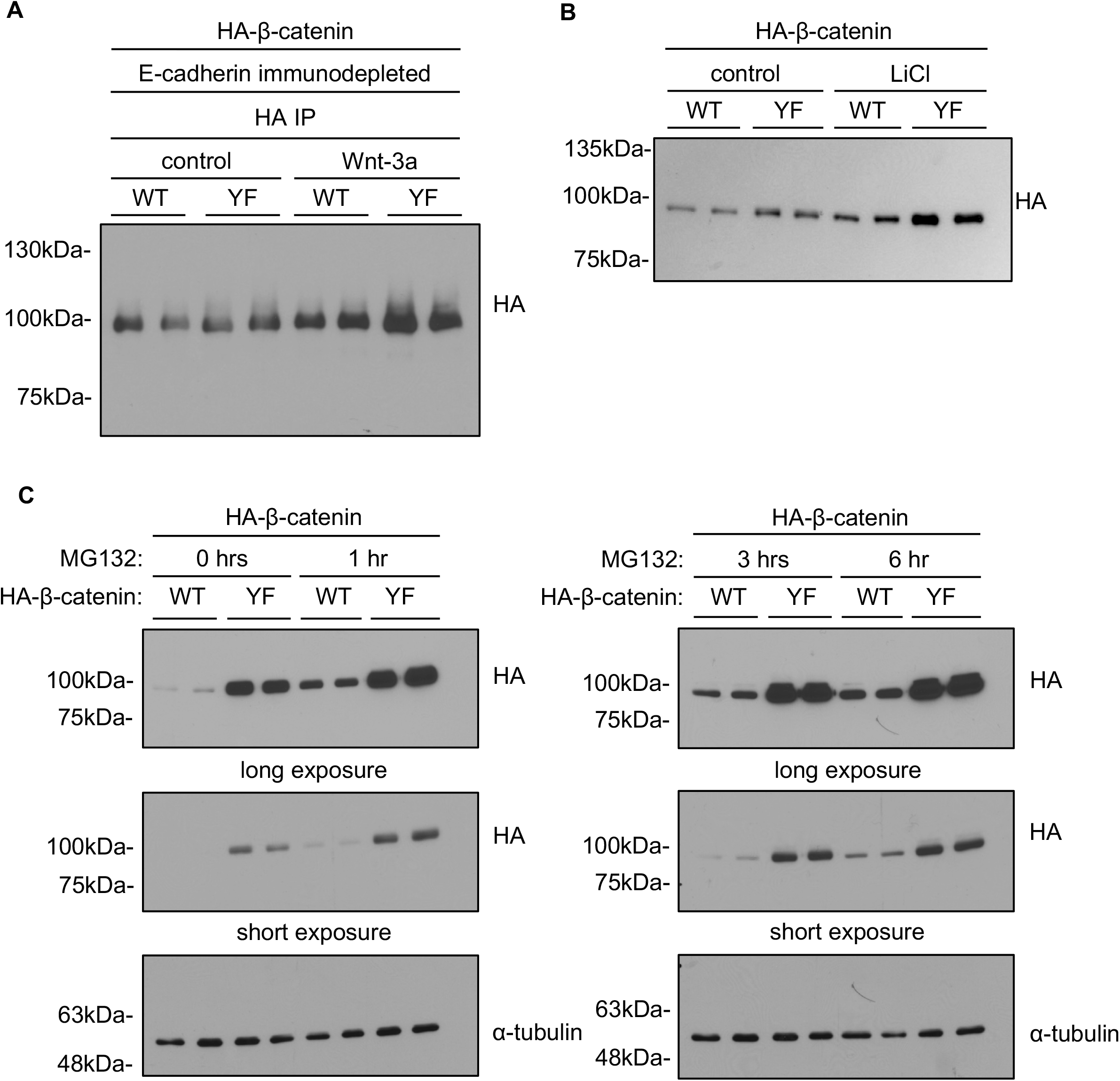
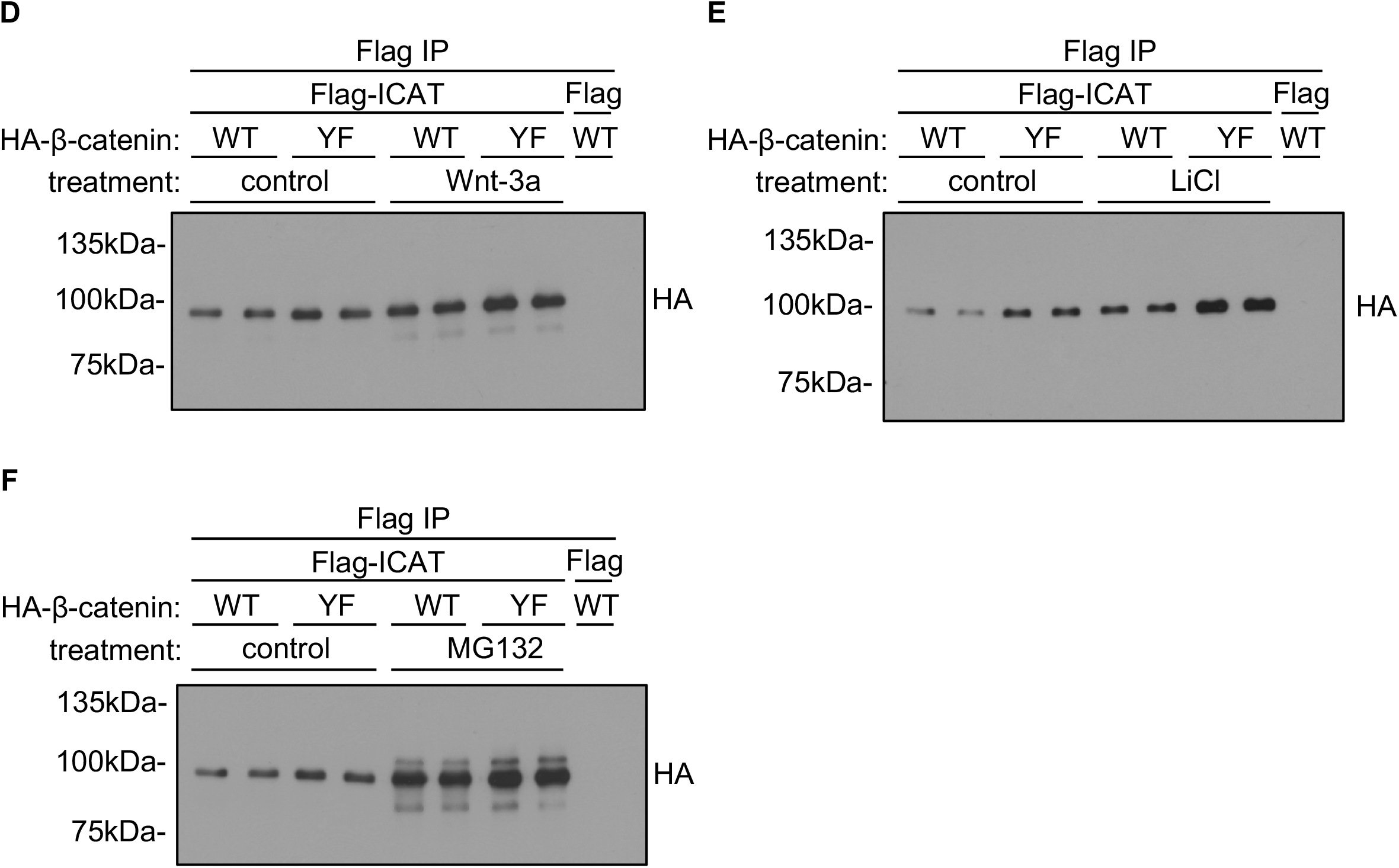
The stabilizing effect of β-catenin tyrosine 30 mutation is independent of the stabilizing effects of Wnt-3a stimulation, GSK3 or proteosomal inhibition. (A) Levels of HA-tagged wild-type (WT) or mutant (Y30F) β-catenin in HEK293T cells treated for 6h with either control or Wnt-3a conditioned media. To focus on the Wnt-responsive β-catenin pool exclusively, lysates were immunodepleted with E-cadherin antibody and concentrated by HA immunoprecipitation prior to analysis. (B) Levels of HA-tagged wild-type (WT) or mutant (Y30F) β-catenin in HEK293T cells left untreated (UN) or grown in the presence of LiCl (GSK3 inhibitor) for 6h. (C) Levels of HA-tagged wild-type (WT) and mutant (Y30F) β-catenin in HEK293T cells grown for 0, 1, 3, and 6h in the presence of the proteosomal inhibitor (MG132). (D-F) Effect of Wnt-3a, LiCl, and MG132 on the levels of exogeneous β-catenin in the (ICAT-associated) signaling pool was evaluated. Levels of HA-tagged wild-type (WT) or mutant (YF) β-catenin co-immunoprecipitating with Flag-tagged ICAT in HEK293 cells grown untreated (control) or in the presence of Wnt-3a-conditioned media for 6hrs (D), LiCl for 6hrs (E) or MG132 for 4hrs (F).

### Wild-type, but not kinase-dead, Nek10 reduces β-catenin association with the Axin complex

Immunoprecipitation experiments revealed a detectable, if weak, interaction between Nek10 and Axin, but not β-catenin (Fig.5A-B). Intriguingly, overexpression of wild-type Nek10 in HEK293T cells caused a considerable reduction in the amount of Axin-associated β-catenin (Fig.5C), consistent with the increased Y30F β-catenin found in Axin immunoprecipitates (see Fig.3D). Expression of the kinase-dead Nek10 mutant failed to produce such an effect, further implying that the Nek10 input into β-catenin turnover is mediated by phosphorylation, rather than another mechanism.

**Figure 5.**
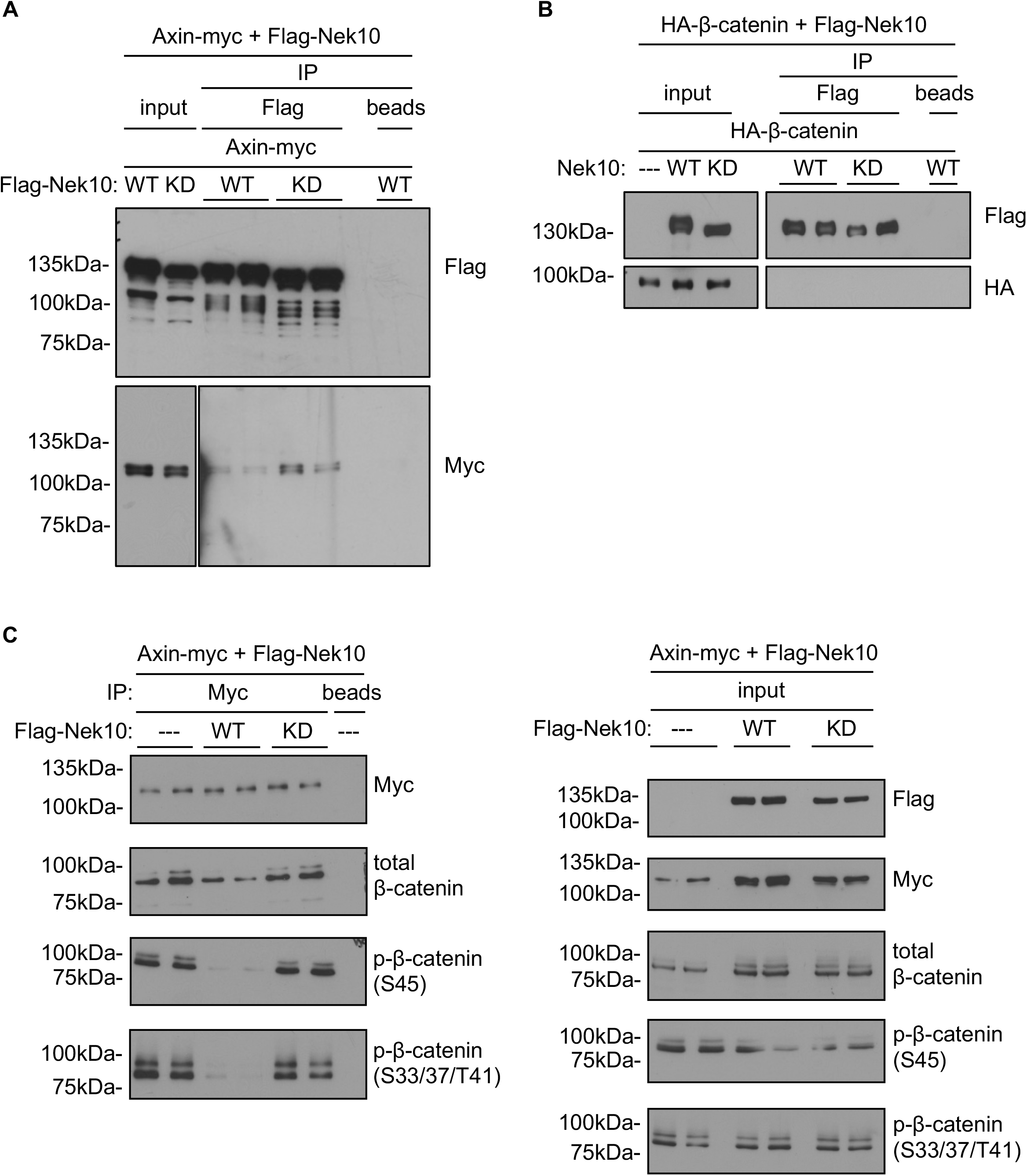
Overexpression of wild-type but not kinase-dead Nek10 results in reduced β-catenin association with the Axin complex. (A) Myc-tagged Axin co-immunoprecipitating with wild-type (WT) and kinase-dead (KD) Flag-tagged Nek10. (B). The ability of wild-type (WT) and kinase-dead (KD) Flag-tagged Nek10 to co-immunoprecipitate HA-tagged wild-type β-catenin was also evaluated. (C) Effect of exogeneous overexpression of wild-type (WT) and kinase-dead (KD) Nek10 on the levels of endogeneous total and Ser45 and Ser33/Ser37/Thr41 phosphorylated β-catenin co-immunoprecipating with Myc-tagged Axin (left panel). Protein levels of respective proteins were determined in whole cell lysates (right panel).

### Effect of Nek10 loss on A549 cell function

To examine the potential physiological effects of Nek10 deficiency, we evaluated *NEK10*^*+/+*^ and *NEK10*^*Δ/Δ*^ A549 cell function in a series of *in vitro* and *in vivo* assays. In suspension tumorsphere forming assays, a measure of stemness in cultured cells, A549 *NEK10*^*Δ/Δ*^ cells were impaired compared to their wild-type controls (Fig.6A). In soft agar growth assays, a measure of tumorigenicity *in vitro, NEK10*^*Δ/Δ*^ A549 cells displayed decreased growth indicating that *NEK10* deletion compromised A549 cell transformation (Fig.6B). Finally, upon tail vein injection into NOD/SCID mice, *NEK10*^*Δ/Δ*^ A549 lines exhibited reduced lung tissue colonization several weeks post injection, compared to controls, measured as cumulative tumor burden (Fig.6C).

**Figure 6.**
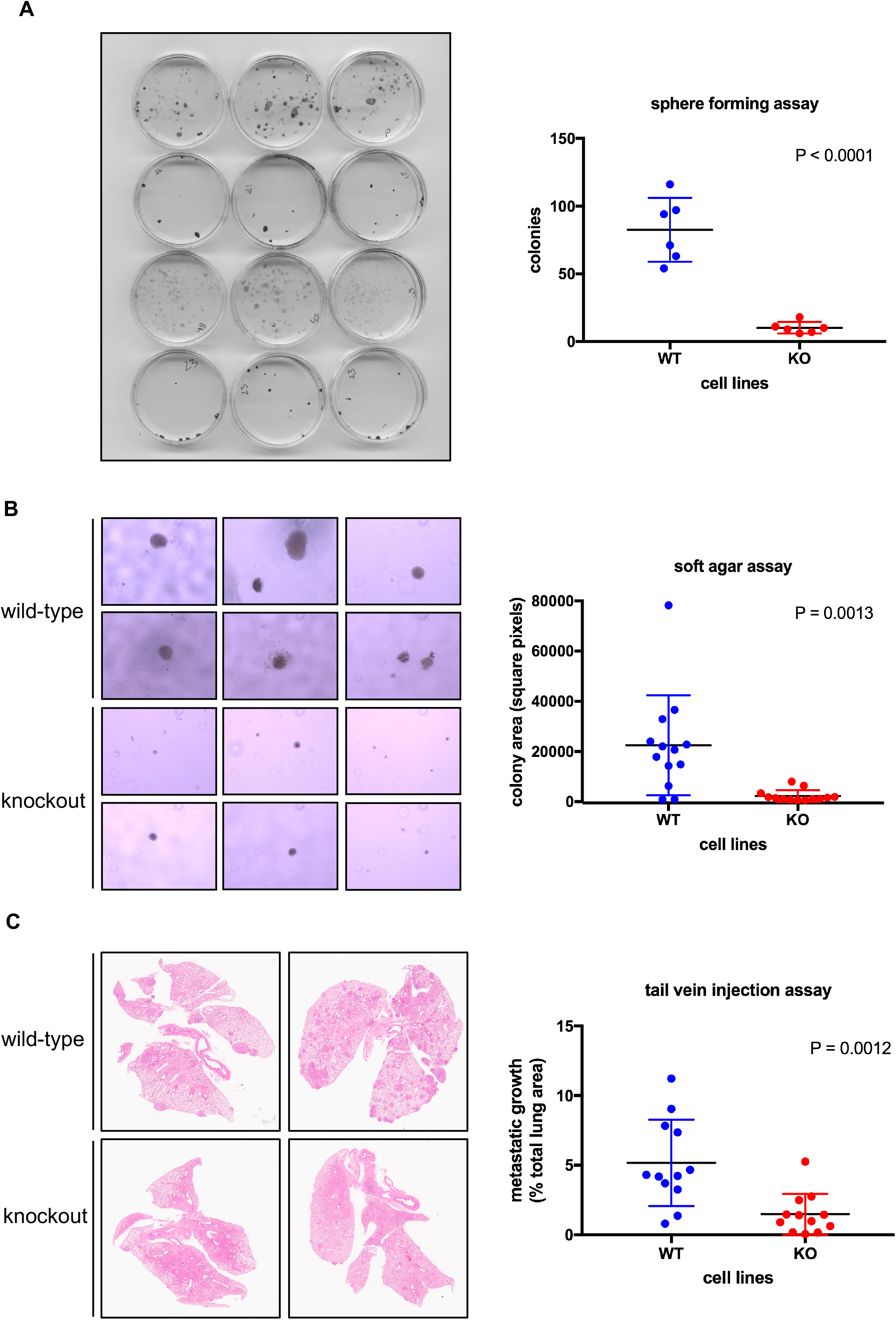
Effect of Nek10 loss on A549 cell function. (A) The capacity of *NEK10*^*+/+*^ and *NEK10*^*Δ/Δ*^ A549 cells to form tumorspheres in suspension was evaluated (unpaired t-test, n=6, error bars represent standard deviation). (B) Anchorage-independent growth of *NEK10*^*+/+*^ and *NEK10*^*Δ/Δ*^ A549 lines was assessed in soft agar assays. Images of four random fields per cell line were analyzed for total collective area of colony growth (unpaired t-test, n=6, error bars represent standard deviation). (C) *In vivo* tumorigenicity was characterized by determining the ability of *NEK10*^*+/+*^ and *NEK10*^*Δ/Δ*^ cells to colonize NOD/SCID mouse lung tissue following tail vein injections. Metastatic growth was quantified by semi-automated digital analysis (unpaired t-test, n=8, error bars represent standard deviation).

## Discussion

Nek10 belongs to the family of NIMA-related cell cycle kinases, but unlike the other members which are serine-threonine kinases, it exhibits a preference for phosphorylation of tyrosines. Comparatively little is known about its cellular substrates and functions. We have previously reported a role for Nek10 in regulating a MAPK-activated G2/M checkpoint in response to UV irradiation (Moniz et al., 2011b). Since it is relatively highly expressed in the lung, we characterized the effect of Nek10 deletion in the A549 lung adenocarcinoma line, leading us to identify a novel function in modulating p53 following DNA damage (Haider et al., 2020). A dramatic upregulation in the baseline levels of β-catenin in both the signaling and adherens junctions pools of the same cells led to the realization of a relationship between Nek10 and β-catenin described here (Fig.1, S3A).

Mechanistically, Nek10 associates with the Axin complex and phosphorylates β-catenin at Tyr30, proximal to the well characterized GSK3 phosphorylation sites at Ser33, Ser37, and Thr41 (Fig.2C-E). Previous work has shown that Tyr30 is one of three β-catenin tyrosines phosphorylated by Jak3, contingent upon prior phosphorylation of Tyr654, but Tyr30 was not further investigated in isolation (Mishra et al., 2017). Phosphorylation of β-catenin could be partially blocked by mutating Tyr30 suggesting that it represents a bona fide Nek10 target site *in vitro* (Fig.2E). It should also be noted that the *in vitro* kinase reactions were carried out in the absence of Axin, which might serve as a scaffold for this process within cells and affect the dynamics of the kinase reaction.

Collectively, Ser33 and Ser37 phosphorylation create a binding site for the β-Trcp E3 ubiquitin ligase leading to the ubiquitination of Lys19 and Lys49 and proteosomal degradation of β-catenin. Upon Ser33/37 phosphorylation, a peptide encompassing residues 17-48 was shown to undergo a significant structural realignment (Megy et al., 2005). Whereas the unphosphorylated peptide is relatively unstructured, Ser33/Ser37 phosphorylation creates a large bend centred on the DpS^33^ GXXpS^37^ β-Trcp binding motif flanked by short helical segments. CK1-phosphorylation of β-catenin at Ser45 primes the GSK3 phosphorylation events by effectively substituting a missing activation loop threonine phosphorylation that is characteristic of many kinases but not GSK3 (Doble and Woodgett, 2003). Nek10 phosphorylation, while not absolutely required, does appear to promote GSK3 phosphorylation as evidenced by the reduction in the relative levels of GSK3-phosphorylated β-catenin in Nek10-deficient cells (Fig.2B). In our studies, CK1- and GSK3-phosphorylated β-catenin are scarcely detectable in cycling A549 cells, but upon treatment with MG132 to arrest proteosomal degradation, the phosphorylated species build up. As there is more free β-catenin available to enter the Axin complex in *NEK10*^*Δ/Δ*^ cells, the levels of CK1-phosphorylated β-catenin are correspondingly elevated (Fig.2B). By contrast, GSK3-phosphorylated β-catenin is actually lower in the Nek10-deficient lines despite the greater substrate availability (Fig.2B). This suggests a potential role for Nek10 in promoting the GSK-mediated phosphorylation steps of the β-catenin turnover cycle. Total cellular GSK3 kinase activity is unaffected by the loss of Nek10 implying that its effect is specific to GSK3 activity within the Axin complex (Fig.S5).

While an association between Nek10 and Axin was observed upon their purification from cells, no direct interaction between Nek10 and β-catenin could be detected (Fig.5A-B). This is consistent with a kinase-substrate relationship, which would favor transient interactions often not detectable by immunoprecipitation. GSK3 has been reported to be constitutively associated with the Axin complex (Li et al., 2012). Thus, it appears that Nek10 does not impact recruitment of either GSK3 or β-catenin to the degradation complex, but is rather constrained to phosphorylation of Tyr30 which presumably confers a kinetic advantage to the GSK3-mediated kinase reaction, possibly via a conformational change.

Blocking phosphorylation of Nek10’s target site in β-catenin (Y30F) led to stabilization of the expressed mutant protein, phenocopying the effect of Nek10 deletion in A549 cells (Fig.3A, S6C). Moreover, relative GSK3-phosphorylation of the mutant β-catenin was also reduced, mimicking the observations in A549 cells (Fig.3B). This in turn led to a relative reduction in the recruitment of the β-Trcp Ubiquitin E3 ligase, providing an explanation for the elevated levels of the mutant protein (Fig.3C). A kinetic assessment of Wnt signaling throughput found that the efficiency of the Axin-mediated turnover is inversely correlated with the relative amount of GSK3-phosphorylated β-catenin (Hernandez et al., 2012). Under conditions of Wnt signaling suboptimal degradation causes an accumulation of β-catenin until a new equilibrium is established, balancing the greater substrate availability with the impaired efficiency of β-catenin turnover, thereby preventing an indefinite rise in β-catenin levels. The absence of Nek10 phosphorylation appears to drive a similar, albeit constitutive, equilibrium. Indeed, compared with the wild-type protein, more Y30F mutant (total and phosphorylated) β-catenin was physically associated with Axin at a given point in time, consistent with the difference in the total cellular levels between the two (Fig.3D). It should be noted that under conditions of Wnt signaling, GSK3-phosphorylated β-catenin has been found to accumulate in both Axin-bound and Axin-free cytosolic pools, revealing that at least a portion of phosphorylated β-catenin can dissociate from the Axin complex (Gerlach et al., 2014). This might explain why impairment in GSK3-mediated phosphorylation of the mutant protein would be more readily apparent at a cellular level than in the specifically Axin-associated pool of β-catenin.

Exogenously expressed wild-type and mutant β-catenin were appropriately localized to both the signaling and adherens junction pools, with mutant β-catenin levels elevated in both (Fig.S6A-B). Moreover, both the wild-type and mutant protein responded as expected to Wnt-3a, LiCl, and MG132 treatments (Fig.4). The stabilization of mutant β-catenin by MG132 implies that the mutant protein is actively degraded by the proteosome and that, in the absence of active degradation, complete saturation of the Axin complex is likely prevented by dissociation of at least some portion of β-catenin. Interestingly, the mutant β-catenin stabilized at higher levels in the presence of Wnt-3a, LiCl, and MG132, suggesting that the effect of Nek10 phosphorylation is independent of those treatments thereby creating an additive effect.

Nek10 regulates p53-dependent transcription by directly phosphorylating p53 at Tyr327 (Haider et al., 2020). Blocking this phosphorylation either through Nek10 deletion or Y327F mutation engenders p53 hypomorphism marked by attenuated transcription of p53 target genes, including p21 and MDM2. This effect is particularly acute following DNA damage and, as a result, Nek10-deficient A549 cells exhibit heightened sensitivity to genotoxic agents. More recently, Axin1 was identified as a potential co-factor with p53 in an *in vivo* Crispr screen in p53+/-mice undertaken to identify genes capable of driving mammary tumorigenesis in the context of p53 heterozygosity (Heitink et al., 2022). Deletion of *Axin1* in p53+/-mouse mammary organoids significantly enhanced the proliferation and altered the morphology of the organoids. Notably, this effect did not appear to be contingent on activation of a Wnt signaling program. Similarly, GSK3 inhibition in Brca1-or Brca2-deficient H1299 cells led to β-catenin accumulation and cell death (Dagg et al., 2021). Mechanistically, this was found to involve β-catenin-dependent downregulation of p21. These interactions between p53 and β-catenin signaling are reminiscent of the phenotypes we observe in Nek10-deficient A549 cells and warrant further study.

β-catenin signaling has a well-established role in maintaining stem cell renewal impacting both normal tissue homeostasis and tumorigenesis (Steinhart and Angers, 2018; Pond et al., 2020). Elevated levels of β-catenin in Nek10-deficient A549 lines correlated with impaired growth of tumorspheres in suspension (Fig.6A). A similar phenomenon was observed in mammospheres grown from mouse mammary epithelial cells, wherein β-catenin stabilization was achieved through the excision of exon 3 encoding the region containing the regulatory GSK3 phosphorylation sites (Dembowy et al., 2015). The upregulation of β-catenin correlated with increased stemness, marked by expansion of a CD24^hi^/CD49^hi^ population, upon exon 3 deletion. Nek10 deletion also impaired the ability of A549 cells to grow in soft agar and colonize the lungs of NOD/SCID mice following tail vein injections (Fig.6B-C). This could reflect changes in stem cell capacity but might also be a consequence of the elevated E-cadherin levels also observed in the Nek10-null lines. Since β-catenin stabilizes E-cadherin by protecting it from endocytosis and degradation, elevated β-catenin and E-cadherin frequently co-occur in lung tumors and more generally (Sormunen et al., 2007). E-cadherin has a well-established tumor suppressive role countering different stages of oncogenesis and its downregulation is a common feature during cancer progression (Chen and Waterman, 2015). For example, heterozygous introduction of the stabilized β-catenin allele promotes tumorigenesis in the small intestine, but not the colon, where E-cadherin levels are naturally higher, without concomitant deletion of E-cadherin (Huels et al., 2015). Collectively, these results suggest that Nek10 loss might impair lung tumorigenesis and are consistent with the observed correlation between reduced Nek10 levels and improved clinical outcome in human lung cancer (Fig.S2).

In summary, we have established a novel role for Nek10 in the regulation of β-catenin levels affecting both its signaling and adherens junctions pools. Nek10 associates with the Axin complex in cells and phosphorylates β-catenin at Tyr30 located proximal to the regulatory sites responsible for controlling β-catenin turnover. In the absence of Nek10, the GSK3-mediated phosphorylation of β-catenin, a step known to be critical for modulating cellular β-catenin levels, was noticeably impaired. Thus, Nek10 phosphorylation of β-catenin at Tyr30, though not strictly required, enhances GSK3 phosphorylation of β-catenin and consequently its ubiquitination and degradation. Nek10 deletion also correlated with reduced tumorigenic potential in several cellular and *in vivo* assays, consistent with current clinical data, possibly implicating Nek10 status as a prognostic marker and therapeutic target in lung as well as other cancers.

### Experimental Procedures

#### Antibodies

The following antibodies were used for Western blotting, immunoprecipitations, and immunofluorescence: total β-catenin (1:2000) (Cell Signaling #9562), phospho-β-catenin S33/S37/T41 (1:1000) (Cell Signaling #9561), phospho-β-catenin S45 (1:1000) (Cell Signaling #9564), α-tubulin (1:2000) (Cell Signaling #2144), PARP (1:1000) (Cell Signaling #9542), pY99 (1:500) (Santa Cruz sc-7020), HA (1:1000) (Cell Signaling #3724), Myc (1:1000) (Cell Signaling #2276), Flag (1:100,000) (Sigma #F3165), GST (1:1000) (Cytiva 27-4577-01), E-cadherin (1:1000) (BD #610181), E-cadherin (Proteintech #20874-1-AP), GAPDH (1:2000) (Santa Cruz #sc-25778), phospho-Lrp6 S1490 (1:1000) (Cell Signaling #2568), phospho-Tau S202/T205 (1:1000)(Cell Signaling #30505). Where applicable, Western blotting dilutions are denoted in parentheses.

### Constructs and Cloning

Generation of the Flag-Nek10 constructs has been described elsewhere (Moniz et al., 2011b). Axin1 was PCR amplified from HT29 cDNA and cloned as a EcoRI/XbaI fragment into pcDNA3.1-myc(C-terminal). β-catenin was amplified from MCF10A cDNA and cloned initially as a KpnI/ApaI fragment into pcDNA3.1-myc(C-terminal), then transferred into pcDNA3.1-HA(N-terminal) as a NotI/ApaI fragment and as a BamHI/NotI fragment into pGEX-4T3 for bacterial expression. The Y30F point mutation was introduced using the QuickChange site-directed mutagenesis kit (Stratagene). GST-ICAT was generated by amplifying a BamHI/XhoI fragment containing ICAT from BT474 cDNA and cloning into pGEX-4T3. The ICAT cDNA was then cloned as a HindIII/EcoRI fragment into p3XFlag-CMV for mammalian expression. Flag-β-Trcp was amplified from A549 cDNA and cloned into p3XFlag-CMV as a NotI/BamHI fragment. Flag-Tau was amplified from T47D cDNA and cloned into p3XFlag-CMV as a NotI-BglII fragment.

### Cell Culture

The generation of the Nek10-KO A549 cell lines has been described previously (Haider et al., 2020). A549, HEK293T, as well as control and Wnt-3a-producing L cells were maintained in DMEM supplemented with 10% fetal bovine serum. Control- and Wnt-3a-conditioned media were generated by passaging L cells at a 1:20 dilution and collecting the supernatant after 96h which was then passed through a 0.45μm filter. When required, unless otherwise specified, cells were treated with 10μM MG132 for 4h, 30mM LiCl for 6h, control- and Wnt-3a-conditioned media diluted with an equal volume of regular growth media for 6h.

### Cell Lysis and Immunoprecipitation

Cells were disrupted in lysis buffer containing 50mM Tris (pH7.5), 150mM NaCl, 1mM EDTA, 1mM EGTA, 1% Triton, 1mM DTT, 1mM sodium orthovanadate, 10mM sodium glycerophosphate, 20mM sodium fluoride, and protease inhibitor cocktail (Roche). Immunoprecipitations were generally carried out with 1mg total lysate incubated with 1μg of primary antibody overnight followed by 90 minutes with Protein A or Protein G sepharose beads and eluted with 2X SDS sample loading buffer. In order to specifically assess exogeneous HA-tagged β-catenin association with Axin, without interference from the endogeneous protein, a two-step immunoprecipitation was carried out as follows. Myc-tagged Axin was immunoprecipitated from 5mg lysate with 4μg of anti-Myc antibody in a volume of 2.5mL overnight, followed by 90 minutes with Protein G sepharose beads. Washed beads were eluted for 15 minutes with 100μL 0.2M glycine pH2.6 supplemented with phosphatase and protease inhibitors. The eluates were first neutralized with 50mM Tris pH8.0, then diluted tenfold for the secondary immunoprecipitation with 1μg anti-HA overnight followed by 90 minutes with Protein A sepharose beads. Immunoprecipitations undertaken to assess β-catenin phosphorylation or the association between β-catenin and β-Trcp were carried out on cells treated for 3hrs with 10 μM MG132 to enable detection. Where required, Western blots were analyzed by Image J (Schneider et al., 2012).

### Subcellular Fractionation

Cell pellets (3×10^6^ cells) were washed with PBS and resuspended in 150μL buffer A (10mM HEPES pH7.9, 10mM KCl, 1.5mM MgCl_2_, 0.34M sucrose, 10% glycerol, 1mM DTT, protease inhibitors). Minimal detergent (0.1% Triton) was gently added to the resuspended cells which were then subjected to a low speed spin at 3,500rpm for 5 minutes following a 7 minute incubation on ice. The supernatant was clarified at 14,000rpm for 5 minutes yielding the cytosolic fraction. The pellet from the low speed spin was washed once in the buffer A, resuspended in buffer B (3mM EDTA, 0.2mM EGTA, 1mM DTT, protease inhibitors) for 15 minutes on ice, then spun at 4,000rpm for 5 minutes with the supernatant retained as the nuclear fraction.

### Recombinant Protein Expression and Purification

Competent BL21(DE3) *E*.*coli* were transformed with the relevant bacterial expression constructs. GST-ICAT was purified from cultures induced at OD_600_ = 0.6 with 0.5mM IPTG for 3 hours at 37°C. Bacterial pellets were disrupted by sonication (5×15sec) in 50mM Tris (pH7.5), 150mM NaCl, 1mM EDTA, 1mM EGTA, 1% Triton, 5% glycerol supplemented with protease inhibitor cocktail (Roche). Recombinant GST-ICAT was batch purified by nutating clarified supernatant with glutathione beads for 2 hours at 4°C, followed by two washes with the above lysis buffer and an additional two washes with a reduced-detergent lysis buffer containing 0.1% Triton. Washed beads were eluted with 50mM Tris (pH8), 150mM NaCl, 1mM EDTA, 5% glycerol, 30mM glutathione for 1 hour at 4°C. Purified GST-ICAT was then dialyzed against 15mM Tris (pH7.5), 300mM NaCl, 5% glycerol for long term storage. Purification of recombinant Nek10 and GST-β-catenin was carried out as described elsewhere (Haider et al., 2020).

### ICAT Pulldown Assay

ICAT pulldown assays were performed by nutating 1mg of cell lysate with 10μg of purified GST-ICAT and glutathione sepharose beads in 1mL total volume for 2 hours at 4°C, followed by three washes with lysis buffer and elution with 2X SDS sample loading buffer.

### Immunofluorescence

A549 cells were grown on coverslips, fixed with 4% paraformaldehyde for 15 minutes at room temperature, permeabilized with PBS/0.1% Triton for 10 minutes at room temperature, blocked with PBS/0.1% Triton/5% normal goat serum (NGS) for 1 hour at room temperature, then incubated overnight at 4°C with the β-catenin primary antibody (1:1000) in PBS/0.1% Triton/5% NGS. The next day coverslips were washed three times with PBS/0.1% Triton and incubated with the anti-rabbit Alexa 488 secondary antibody for 1 hour at room temperature. Coverslips were washed three times with PBS, counterstained with DAPI for nuclear visualization, mounted in Mowiol (Sigma), then analyzed by confocal microscopy.

### Quantitative RT-PCR

Total RNA was purified from cells using the PureLink RNA Miniprep kit (Invitrogen). cDNA was synthesized by reverse transcribing 200ng of total RNA in a volume of 20μL using the SensiFAST kit (Bioline). Quantitative real-time PCR reactions (10μL) were then run in the Roche LightCycler 480 machine using 1μL of the cDNA reactions as template, 200nM forward and reverse primers, and 2X Sybr mix (Bioline SensiFAST Sybr No-Rox kit). The gene-specific primer sequences can be found in Table 1. Each reaction set was performed in triplicate and relative gene expression was calculated using the Livak method (2^-ΔΔCt^) with GAPDH as the reference gene.

**Table 1.**
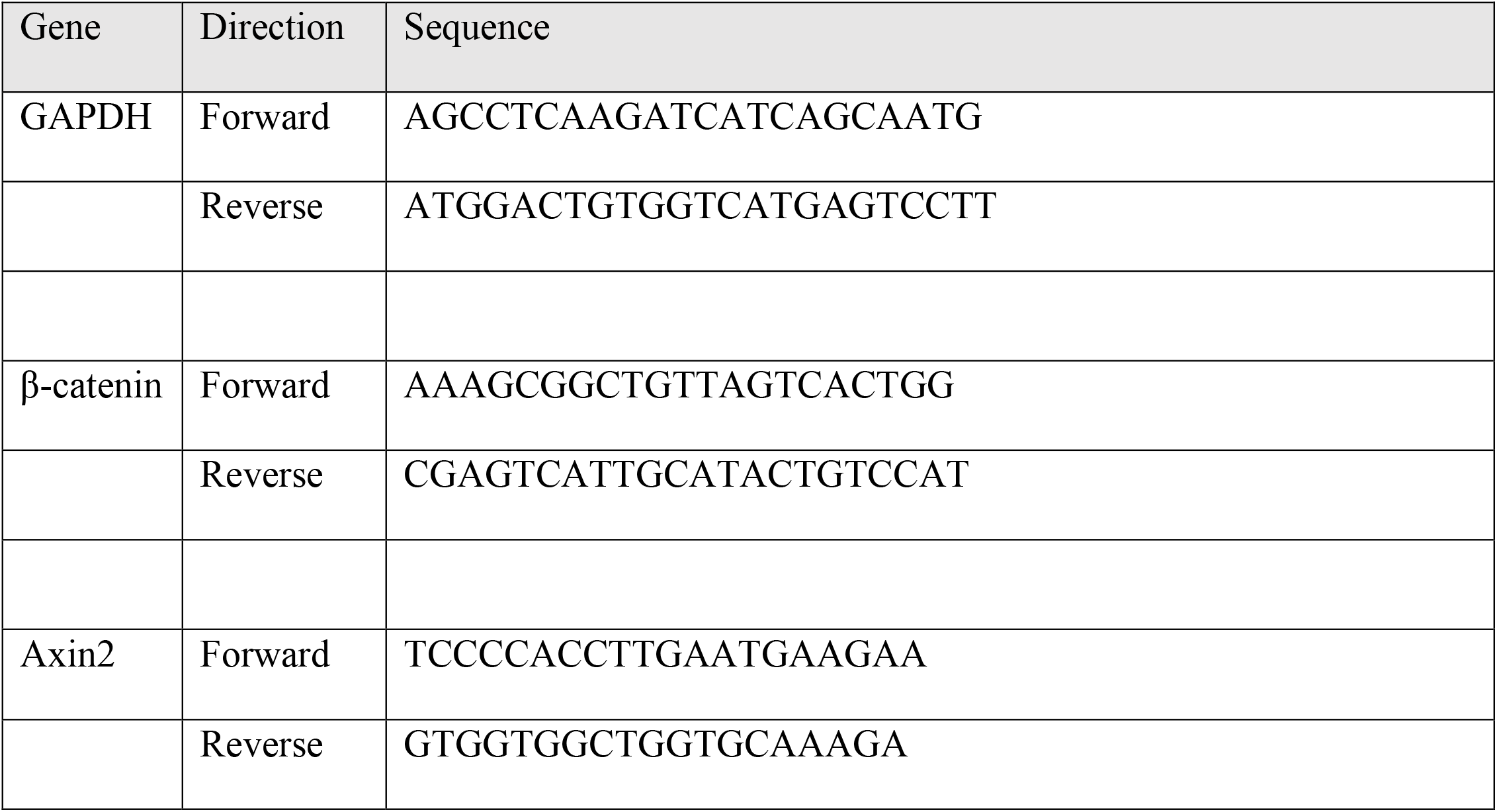
Primers used for quantitative RT-PCR

### Kinase Assays

The ability of Nek10 to in vitro phosphorylate β-catenin was evaluated using both radioactive and non-radioactive assays. In the radioactive assay, purified recombinant Nek10 (wild-type or kinase-dead) was used to phosphorylate β-catenin immunoprecipitated from either *NEK10*^*+/+*^ or *NEK10*^*Δ/Δ*^ A549 cells. Reactions were carried out for 30 minutes at 30°C in 40μL volume containing 50mM MOPS (pH7.4), 10mM MgCl_2_, 10mM MnCl_2_, 2mM EGTA, 20mM sodium glycerophosphate, 1mM DTT, 5uM ATP and 5μCi [γ-^32^P]ATP. Reactions were terminated with 2X SDS sample loading buffer, separated by SDS-PAGE and detected by autoradiography. The non-radioactive assay was carried out similarly using recombinant Nek10 kinase, but with recombinant purified GST or GST-β-catenin (wild-type or Y30F mutant) as the substrate, and omitting [γ-^32^P]ATP from the reactions, which were resolved by SDS-PAGE and detected by Western blot with pY99 antibody.

### GSK3 Activity Assay

The levels of GSK3 activity in A549 cells were evaluated by monitoring the levels of exogeneous Tau phosphorylation at Ser202/Thr205. Since endogeneous Tau is expressed sparingly in A549 cells, *NEK10*^*+/+*^ and *NEK10*^*Δ/Δ*^ lines were transfected with a Flag-Tau construct. Exogeneous Tau was immunoprecipitated from cell lysates 48hrs later and Tau phosphorylation in the immunoprecipitates determined using a phospho-Tau antibody.

### Tumorsphere Assay

A549 cells were tryspinized, resuspended in sphere forming media consisting of serum-free DMEM/F12 supplemented with B27 (1:50), 20ng/mL EGF, 20ng/mL FGF, and 10μg/mL insulin, then plated at 1000 cells/well in a 6-well plate. After 2 weeks of growth, cultures were spiked with 10% fetal bovine serum overnight to permit attachment for quantification.

### Soft Agar Assay

Assays were carried out in 35mm low attachment plates. A base layer was prepared consisting of 2mL regular growth media supplemented with 0.7% low melt agarose. The top layer had 1000 cells/plate in 1mL regular growth media supplemented with 0.35% low melt agarose. Cells were grown for two weeks with fresh growth media overlayed every four days. Plates were imaged and colony growth quantified using ImageJ.

### Tail Vein Injection Assay

A549 cells were trypsinized, resuspended in phosphate buffered saline (PBS), and 1×10^5^ cells were injected in 100μL into the tail veins of 12-week old NOD/SCID mice. After 6 weeks, mice were euthanized with CO_2_, after which the lungs were dissected, fixed in 10% formalin, sectioned, and stained with haematoxylin and eosin (H&E) for analysis. Lung metastasis area was quantified by semi-automated digital analysis using QuPath pathology and bioimage analysis software (Bankhead et al., 2017). Briefly, digital images of H & E-stained lung sections were loaded into a single project file. The *Simple Tissue Detection* command was first used to batch process the selection of lung tissue and exclusion of white space on all images, followed by manual annotation of each tissue sample and exclusion of staining artifacts. A random trees-based pixel classifier was manually trained to detect normal tissue and metastatic regions and then applied to each annotated lung section. Detections of metastasis made by the classifier were converted to annotation-type objects, and area measurements of each annotation were automatically tabulated. Percent tumour area was then calculated by first totaling the area of all tumour detections within a given lung annotation and dividing this sum by the area of the parent lung annotation.

### Bioinformatics

Nek10 expression in human tissues was evaluated using the GTEx portal. The Genotype-Tissue Expression (GTEx) Project was supported by the Common Fund of the Office of the Director of the National Institutes of Health, and by NCI, NHGRI, NHLBI, NIDA, NIMH, and NINDS. Nek10 expression in mouse tissues was derived from RNAseq data deposited in the NCBI gene expression omnibus (Barrett et al., 2012) from two independent studies, GSE41637 (Merkin et al., 2012) and GSE74747 (Huntley et al., 2016). The correlation between Nek10 expression levels and clinical outcome in patients with lung adenocarcinomas was evaluated using KM-plotter (Lanczky et al., 2021). The analysis was run, exclusively on adenocarcinomas, using a combined data set which including the following cohorts: TCGA, CAARRAY, GSE14814, GSE19188, GSE29013, GSE30219, GSE31210, GSE3141, GSE31908, GSE37745, GSE43580, GSE4573, GSE50081, GSE8894.

---

**Figure S1.**
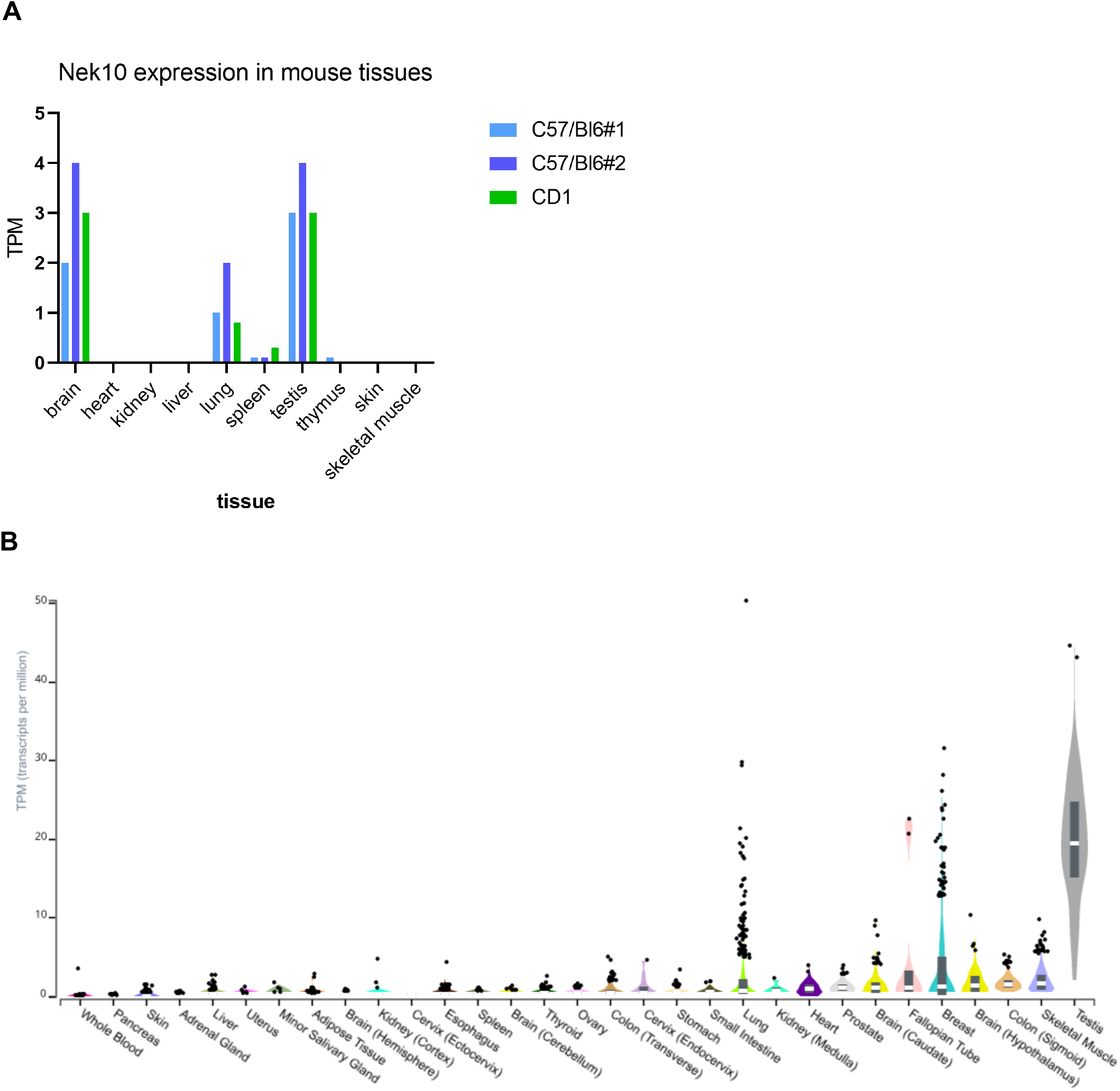
Nek10 tissue expression. (A) Nek10 expression in mouse tissues was assessed using RNAseq data deposited in the NCBI gene expression omnibus from two independent studies. (B) Nek10 expression in human tissues was determined from the Genotype-Tissue Expression (GTEx) project RNAseq database.

**Figure S2.**
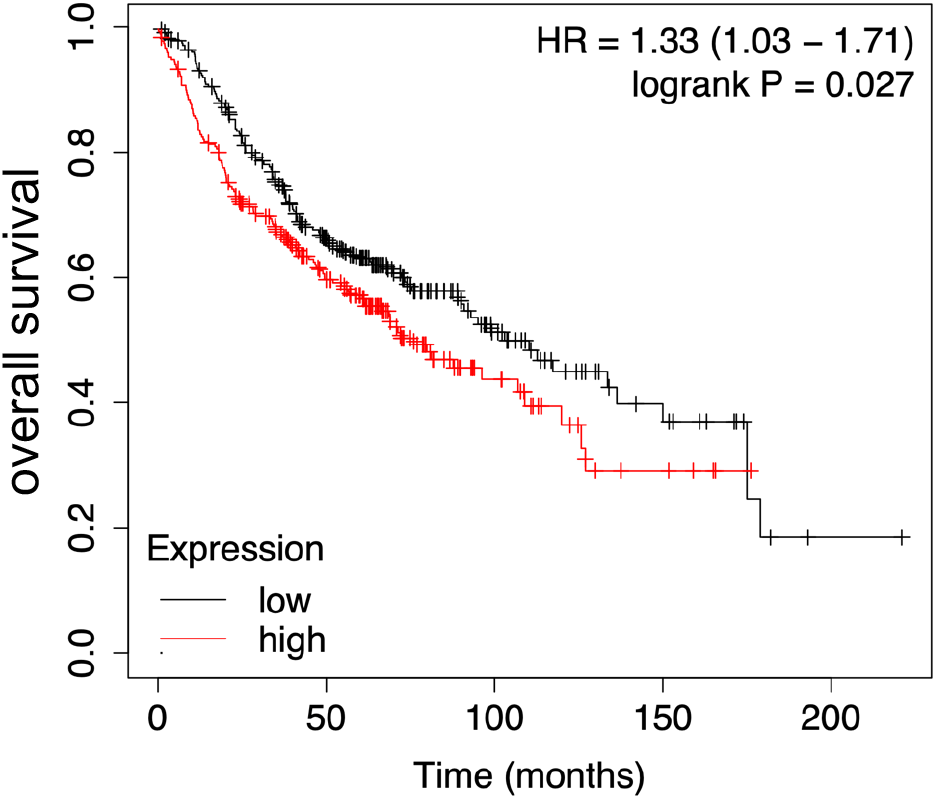
Correlation between lung cancer outcome and Nek10 expression. Kaplan-Meier plot derived from KM-plotter charting the overall survival of patients diagnosed with lung adenocarcinomas stratified by Nek10 expression. Combined data from a number of independent cohorts was used in the analysis, as described in the Methods.

**Figure S3.**
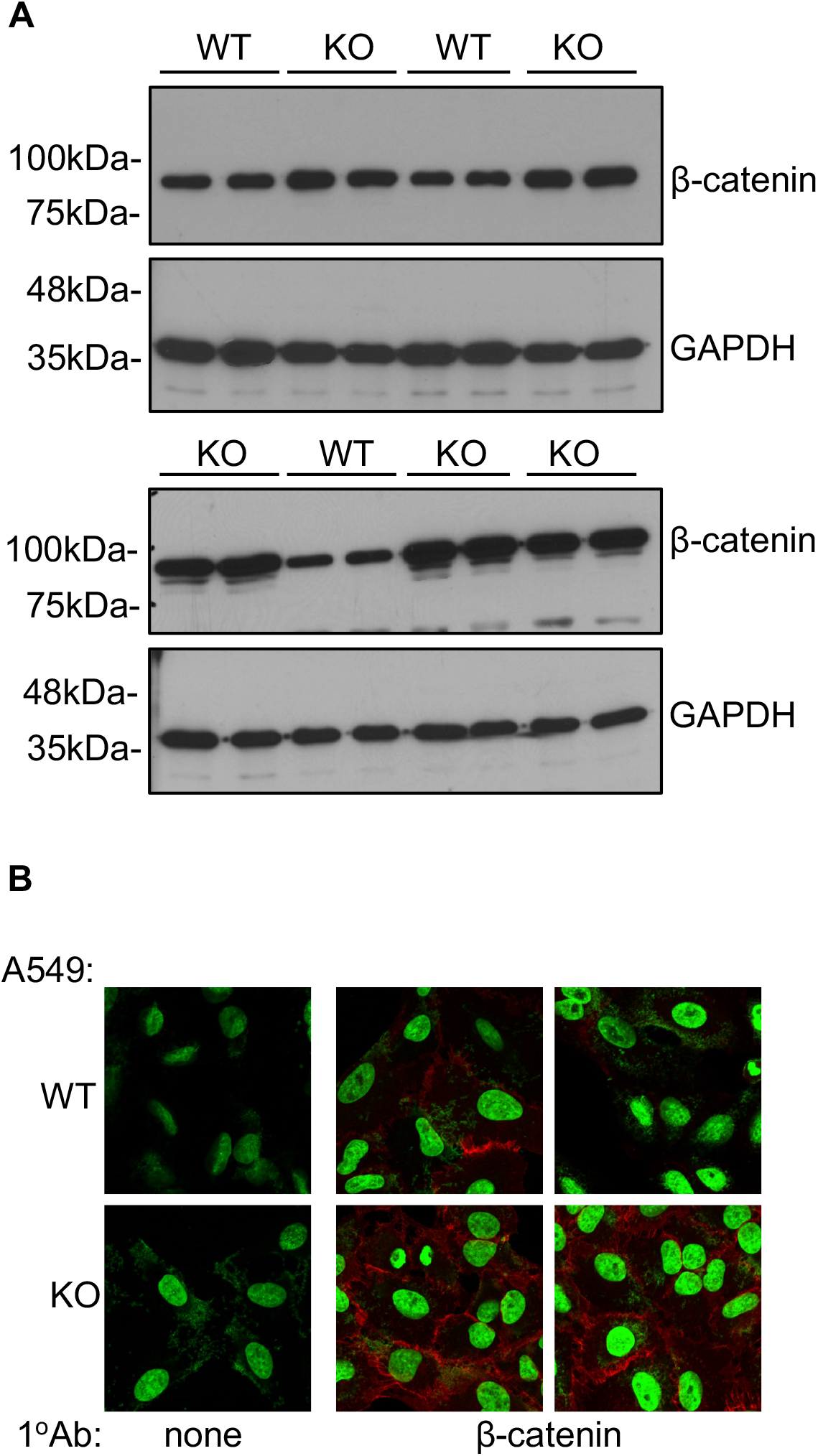
Nek10 deletion results in elevated β-catenin in A549 cells. (A) Total β-catenin levels in whole cell lysates from three *NEK10*^*+/+*^ and five *NEK10*^*Δ/Δ*^ A549 lines. GAPDH was used as a loading control. (B) β-catenin localization in *NEK10*^*+/+*^ and *NEK10*^*Δ/Δ*^ A549 cells assessed by immunofluorescence.

**Figure S4.**
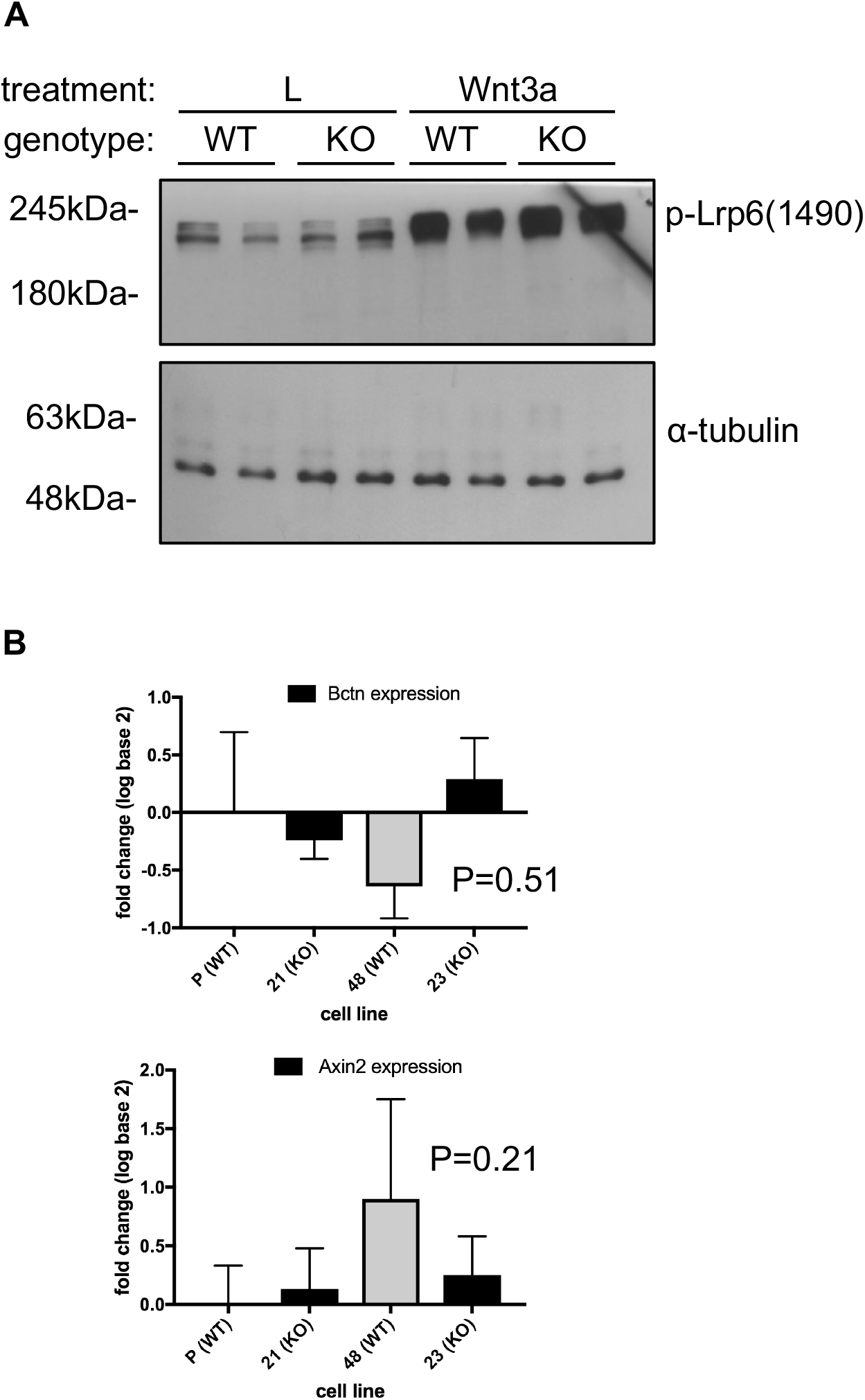
Nek10 deletion does not affect upstream Wnt pathway activation in A549 cells. **(A)** Phosphorylation of the Lrp6 receptor in *NEK10*^*+/+*^ and *NEK10*^*Δ/Δ*^A549 cells after 6hrs treatment with either control or Wnt-3a-conditioned media. α-tubulin was used as a loading control. (B) Relative expression of β-catenin and the β-catenin target, Axin2, in *NEK10*^*+/+*^ and *NEK10*^*Δ/Δ*^ A549 cells determined by quantitative RT-PCR (two-tailed t-test, n=3, bars represent standard deviation).

**Figure S5.**
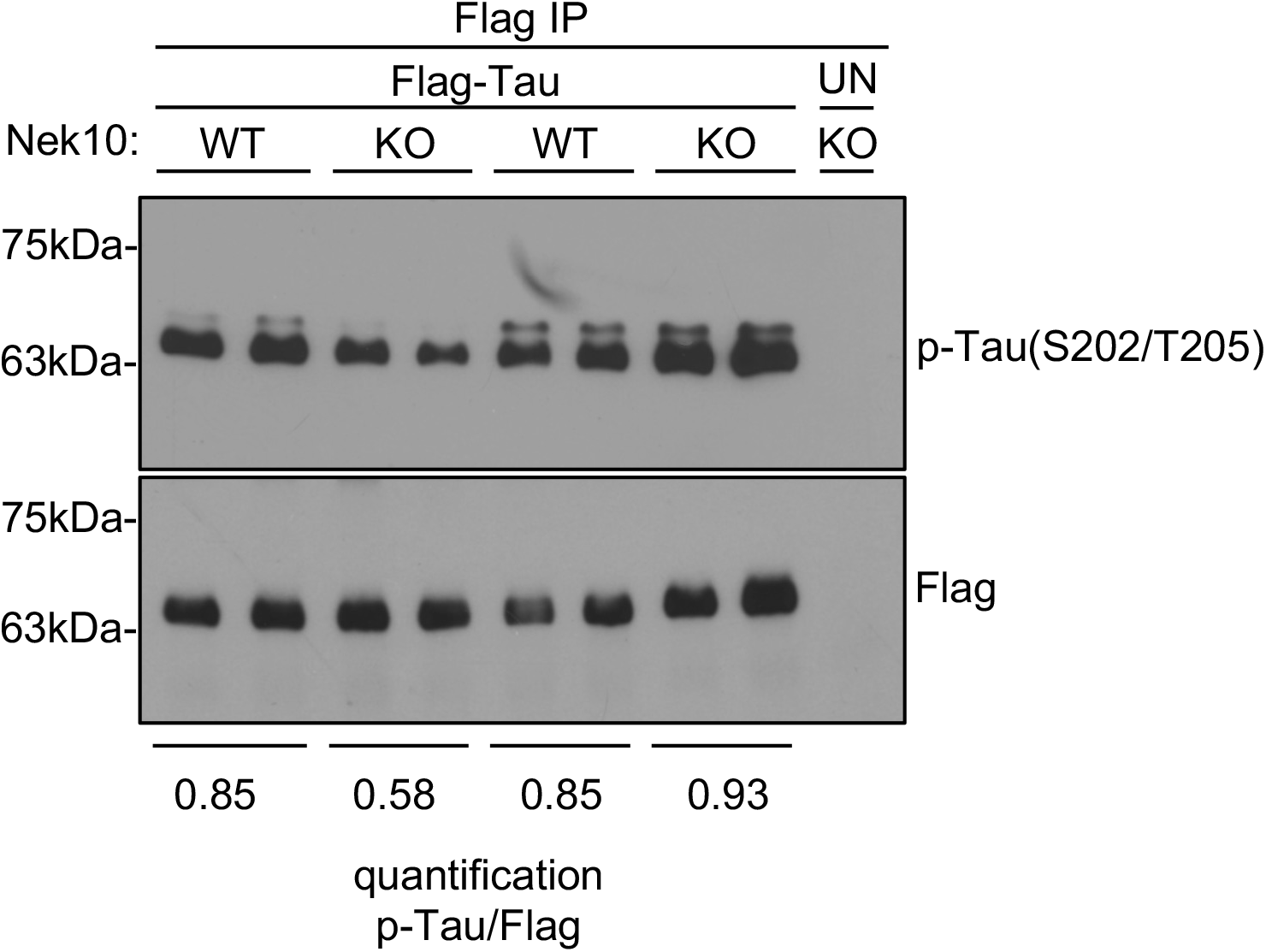
Nek10 deletion does not affect overall GSK3 activity in A549 cells. Tau phosphorylation, a readout of GSK3 activity, was evaluated in *NEK10*^*+/+*^ and *NEK10*^*Δ/Δ*^ A549 cells. ImageJ analysis was used to quantify Tau phosphorylation, normalized to the total Flag-Tau immunoprecipitated.

**Figure S6.**
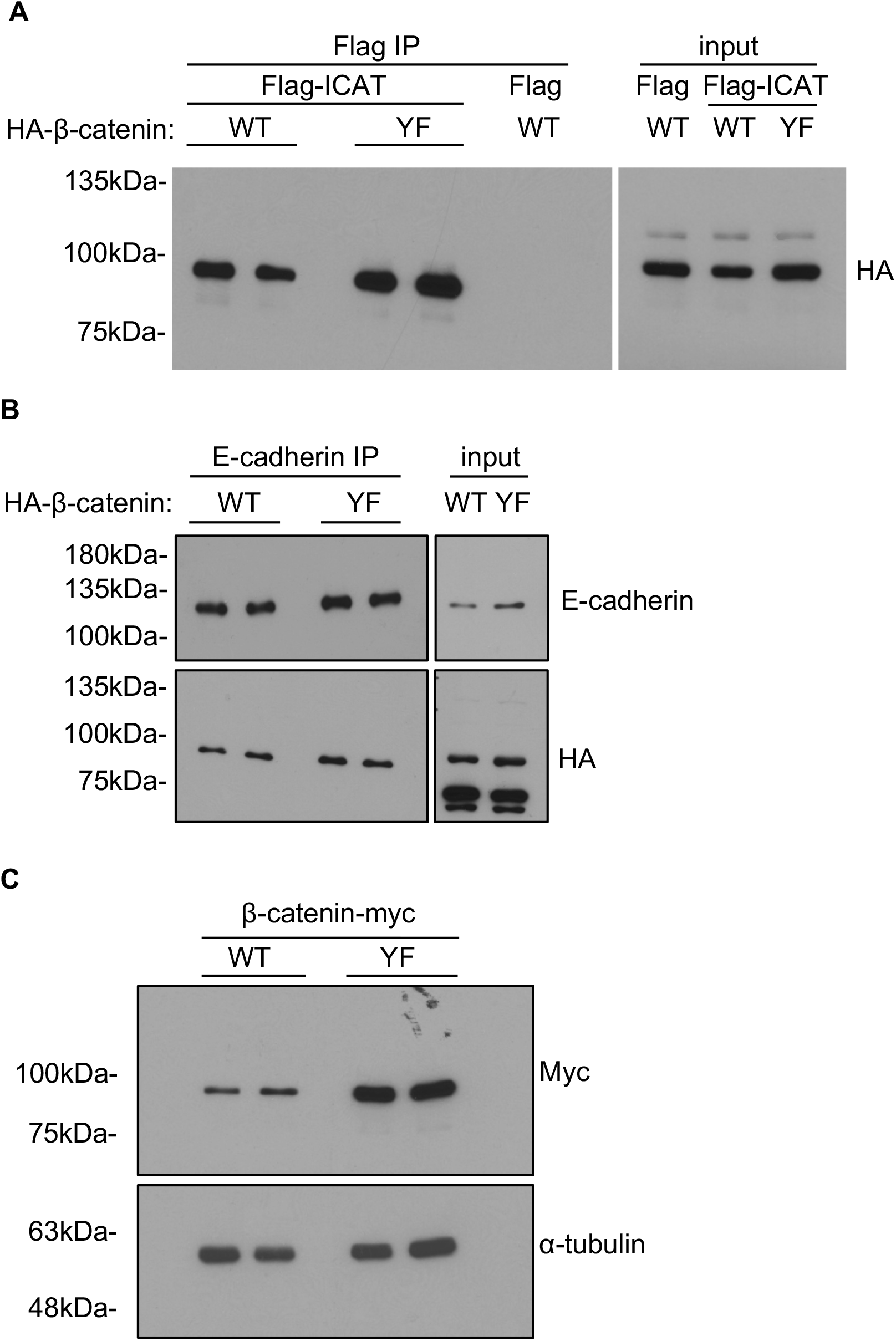
Tyrosine 30 mutation results in increased β-catenin levels. (A-B) Proper localization of HA-tagged β-catenin to both the signaling and adherens junction pools was verified by monitoring the co-immunoprecipitation of HA-tagged wild-type (WT) or mutant (Y30F) β-catenin with either Flag-ICAT (A) or E-cadherin (B). (C) The levels of C-terminal Myc-tagged wild-type (WT) and mutant (Y30F) β-catenin in HEK293T cells. α-tubulin was used as a loading control.

